# Transcriptomic timeseries links hepatic gene expression to an early and self-limited systemic response to enteric infection

**DOI:** 10.64898/2026.04.07.717127

**Authors:** Yuko Hasegawa, Akina Osaki, Masataka Suzuki, Ting Yan, Ian W. Campbell, Matthew K. Waldor

**Author notes:** These authors contributed equally (Yuko Hasegawa, Akina Osaki, Masataka Suzuki).

## Abstract

The liver receives microbe and host signals from the intestine via the portal vein, thereby connecting the gut to systemic physiology. Homeostatic control of the timing of systemic responses is critical to prevent the expansion and dissemination of gut microbes and to mitigate untoward effects from prolonged systemic inflammation, however these mechanisms remain enigmatic. Here, to determine the role of the liver in coordinating systemic immune responses to enteric infection, matched measurements of global gene expression profiles were collected from the murine liver and intestinal epithelium throughout the course of enteric infection and clearance of *Citrobacter rodentium*, a mouse model of infectious colitis. These data revealed metabolic suppression in the liver during the peak of infection and a long-lived immune signaling pattern in the colon associated with CD4 and CD8 T cell infiltration that persisted beyond the clearance of infection. Furthermore, an early inflammatory signal was detected in the liver that resolved before the peak of disease and pathogen colonization. This self-limited, early signal depended on the pathogen’s virulence program and correlated with the timing of a corresponding systemic response, including circulating TNF-α and IL-6, key mediators of acute-phase proteins. These results uncover the temporal pattern of hepatic changes in response to the course of intestinal infection and provide correlative evidence that an early pulse of gene expression in the liver coordinates and limits the duration of the systemic acute-phase protein response.

**Author summary:** Localized infections can trigger systemic inflammatory responses that help limit infection. However, prolonged systemic inflammation risks tissue damage and disease. Thus, the timing of systemic immune reactions is critical to the balance between immunity and damage. Yet, knowledge of the pathways that link gut-localized infections to systemic immune tone is limited. Since the liver is anatomically central to the connection between the gut and circulatory systems, we hypothesized that monitoring the kinetics of liver gene expression throughout the natural course of an intestinal bacterial infection would provide a valuable resource and yield insight into the control of systemic immune tone. Critically, we identified a burst of liver cytokine signaling that occurred and resolved before peak pathogen burden and disease in the colon and predicted the circulating inflammatory response. We propose that this early and self-limited signal from the liver coordinates the timing of the systemic response, ensuring it occurs early enough to promote immunity and resolve before causing tissue damage.

## Introduction

Localized infections often stimulate systemic responses that circulate throughout the body. Such responses include the acute phase response, which changes the abundance of several hundred serum proteins, such as complement components, fibrinogen, and C-reactive protein (CRP), that accompany both localized and disseminated infections[1]. Generally, these responses are self-limited and resolve as the infection clears. However, sustained inflammatory signals, such as IL-6, can lead to tissue damage and disease[2,3]. Indeed, inhibiting IL-6 can help prevent tissue damage caused by prolonged inflammation[4,5]. Further research is needed to deepen the understanding of the signals and mechanisms by which localized infection stimulates and ultimately constrains prolonged systemic inflammation.

The consequences of intestinal infection on systemic immune tone, as well as the pathways, mediators, and mechanisms by which enteric infection modifies systemic responses, remain relatively uncharted. The liver is anatomically linked to the intestine via the portal circulation and to the systemic circulation, connecting these two systems. Besides nutrients from digestion, the portal circulation delivers inflammatory mediators and gut microbe-derived products, including metabolites[6], microbe-associated molecular patterns (such as lipopolysaccharide)[7], and even intact bacteria to the liver[8] (reviewed in [3,9–14]). At least in part in response to this gut microbe-derived input, the liver produces a wide array of immunomodulatory molecules, including acute phase proteins[1,15], metabolites[16], and bile acids[17–19] that are delivered to the blood and, in some cases, to the bile. The liver’s output in bile, which is emptied into the duodenum, can feed back on the gut and impact intestinal and microbial physiology.

Since the liver represents the critical nexus between the intestine and the circulatory system, we hypothesized that charting the kinetics of hepatic global gene expression changes throughout the natural course of an enteric infection could provide insight into the control of systemic responses to intestinal infection. Here, we used *Citrobacter rodentium*, a murine pathogen that serves as a model for the human food-borne pathogens enteropathogenic and enterohemorrhagic *Escherichia coli* (EPEC/EHEC)[20], to generate matched timeseries data comprising pathogen burden and RNA abundance from the liver and the colonic epithelium. These ‘attaching and effacing’ pathogens cause diarrheal disease by attaching to the intestinal epithelium using a conserved type III secretion system that also injects effector proteins that modify various host processes including reducing intestinal barrier function[21–23]. Unexpectedly, these data revealed an early hepatic gene expression program that resolved before peak pathogen burden and disease in the colon. The early, self-limited hepatic response primarily consisted of transcripts associated with inflammatory responses, including IL-6 and TNFα, key activators of the acute phase response[1,24–26]. Notably, corresponding serum protein abundance was strongly correlated with the timing of hepatic gene expression. We propose that the early, transient hepatic response explains the temporal segregation of the acute phase response to an early window during enteric infection.

## Results

### Enteric infection induces an early pattern of gene expression in the liver

To understand the hepatic response to enteric infection, a matched dataset was created to chart pathogen burden and host RNA abundance in the liver and colonic epithelium throughout the course of *C. rodentium* infection in C57BL/6J mice. This extracellular pathogen is a natural agent of murine colitis and is generally considered an intestine-restricted pathogen; however, small numbers of *C. rodentium* have been detected outside the gut in some studies[27,28], likely due to pathogen translocation subsequent to infection-induced damage to gut barrier integrity[29]. Following oral gavage of 5×10^9^ *C. rodentium* colony-forming units (CFU), *C. rodentium* expansion was detected in both the cecum and colon by 3 days post-inoculation (dpi; Fig. 1A, Fig. S1). Colonic colonization peaked 6-10 dpi at ∼10^9^ CFU/g and then declined and became undetectable in most animals by 21 dpi, consistent with previous observations[28,30]. During the peak of intestinal colonization on day 6 and 10, some animals had a small number of detectable *C. rodentium* (∼10^3^ CFU/g) in the liver (Fig 1A). However, the hepatic burden was generally <1/10^6^ of the colonic burden, and hepatic CFU were not observed 1-3 dpi or after 10 dpi.

**Fig. 1.**
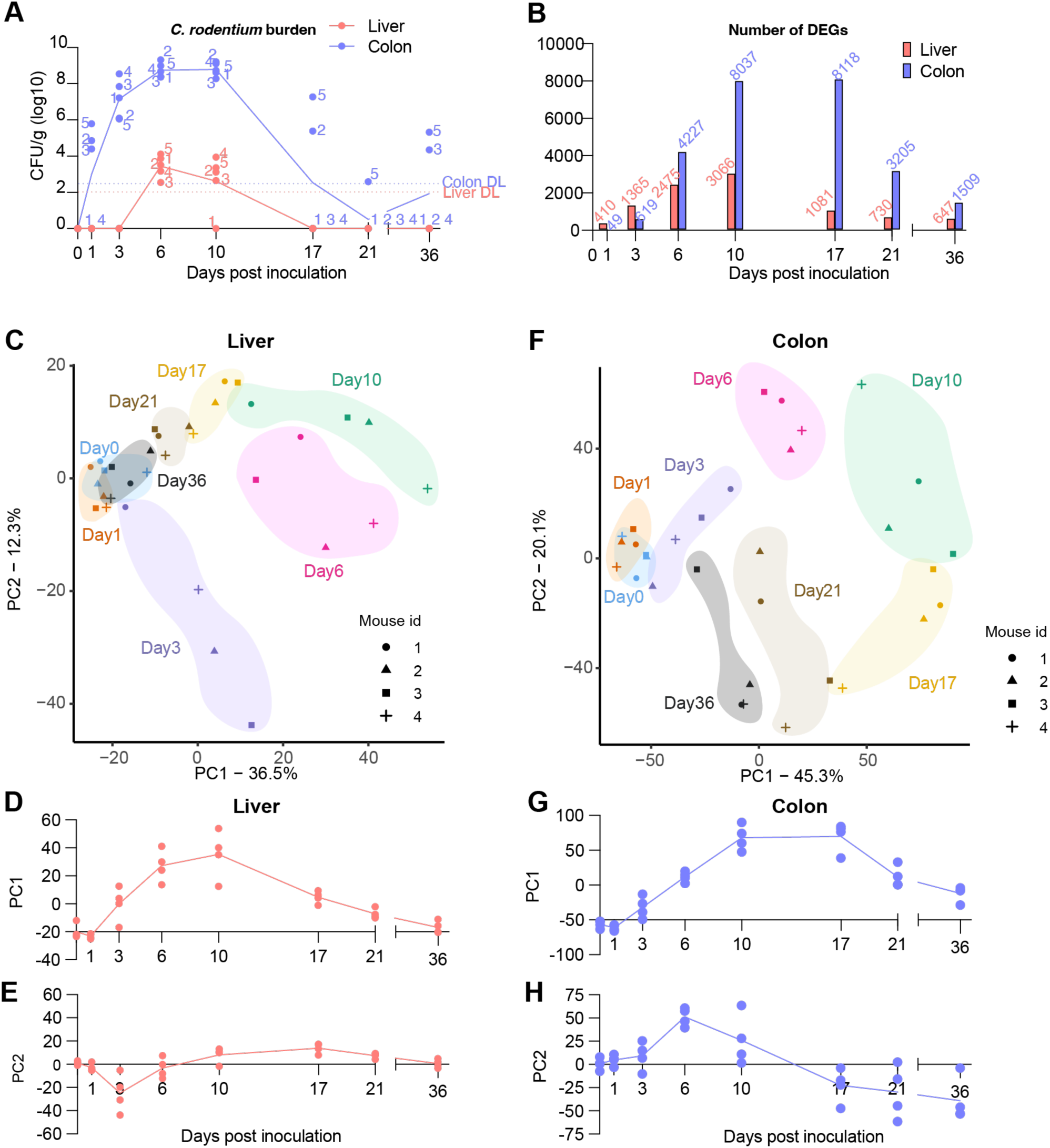
Temporal patterns of liver and colonic epithelial gene expression profiles in response to *C. rodentium* infection. **A.** *C. rodentium* burden (colony-forming unit per gram of tissue: CFU/g) in the liver (red) and the colon (blue) at each time point (days post inoculation) following inoculation with 10^9^ CFU. Points represent mice, and the numbers on the dots indicate mouse ID. Solid lines indicate geometric mean. Horizontal dotted lines indicate the detection limit (DL) in the liver (red), and the colon (blue). **B.** Bar graph showing the number of differentially expressed genes (DEGs) with the cut-off of absolute log_2_ fold change > 1 and adjusted *p*-value < 0.05 in the liver (red) and the colonic epithelium (blue) at each time point (days post inoculation) compared to uninfected day 0. The number of DEGs are shown on top of the bars. **C-H.** PCA plots showing the transcriptome profiles in the liver (C) and the colonic epithelium (F). Colored areas represent the time points (day 0: blue, day 1: orange, day 3, purple, day 6: pink, day 10: green, day 17: yellow, day 21: brown, day 36: gray), and the symbols represent mouse ID. Scatter charts show PC1 and PC2 values (D: liver PC1, E: liver PC2, G: colon PC1, and H: colon PC2) at each time point (Day 0 – Day 36). Mouse IDs among Fig. 1A, C and F are matched.

Unexpectedly, the gene expression patterns associated with enteric infection detected in the liver generally preceded those in the colonic epithelium. On days 1 and 3 post-inoculation, when *C. rodentium* was detected in the colon but not in the liver (Fig. 1A), there was a greater number of differentially expressed genes (DEGs) in the liver vs the colonic epithelium (Fig. 1B, Supplementary Table 1). By contrast, from day 6 onwards, the number of colonic DEGs exceeded those in the liver. The highest number of DEGs in the colonic epithelium was apparent starting 10 dpi, corresponding to peak colonization (Fig. 1B). However, the peak of the colonic expression profile continued to 17 dpi, when the *C. rodentium* burden was declining. Notably, thousands of colonic DEGs remained at 21-36 dpi, a time when most animals had already cleared the pathogen. Thus, this gross tally of the numbers and time course of colonic and hepatic DEGs indicates that the liver responds earlier to enteric infection than the colonic epithelium, but that the colonic epithelium exhibits more complex and longer-lived responses than the liver. These differences in the number and kinetics of DEGs in the liver and colon demonstrates that enteric infection induces temporally distinct transcriptional reprogramming in local and distal organs.

Principal component analyses (PCA) also captured the distinct patterns of gene expression changes in the liver and colon in response to *C. rodentium* infection (Fig. 1C-H, Fig. S2). Overall, in the liver, a substantial response to infection was observed as early as 3 dpi (Fig. 1CDE). The trajectory of liver PC2 was notable because a distinct signal was apparent only on day 3 and reversed toward baseline thereafter (Fig. 1CE); this self-limited change in hepatic expression manifested before the pathogen was detected in the liver (Fig. 1A). By contrast, liver PC1 increased in magnitude following the same pattern as pathogen burden in the colon, peaking at 10 dpi and declining thereafter (Fig. 1CD). Interestingly, hepatic PC1 and PC2 returned to their pre-infection baseline by 36 dpi, overlapping with their day 0 values (Fig. 1CDE), suggesting that the effects of enteric infection on hepatic gene expression are generally short-lived.

As anticipated, infection resulted in marked changes in colonic epithelial gene expression profiles. Like liver PC1, the trajectories of colon PC1 and PC2 initially increased with pathogen burden, rising until day 6-10 (Fig. 1FGH). However, PC1 remained constant from 10-17 dpi, reflecting an ongoing response that persisted as the pathogen burden declined. Furthermore, despite pathogen clearance, the colonic epithelial transcriptome did not return to the day 0 baseline 36 dpi (Fig. 1 FGH), suggesting that the colonic epithelial gene expression program bears a lasting mark of infection.

The distinct kinetics of expression changes in the liver and colon were also apparent when the longitudinal data were partitioned into clusters (Fig. S3). By 3 dpi, changes in gene expression were detectable in both organs; however, the early increase in hepatic cluster 1 gene expression was distinctive, as it was the only pattern to revert after this early time. The peak of hepatic expression changes mainly resolved beyond 10 dpi (liver clusters 1-4), but several changes in colonic gene clusters remained altered through 36 dpi (colon clusters 3, 4, and 6).

### Enteric infection transiently diminishes hepatic expression of metabolic pathways

Gene set enrichment analysis (GSEA)[31,32] using metabolic pathways in KEGG[33] was used to identify the major pathways in the liver and colon whose RNA abundance changed in response to *C. rodentium* infection. These analyses showed that infection was associated with reduced expression of genes linked to diverse metabolic pathways in the liver, but generally not in the colon (Fig. 2A, Supplementary Table 2). Reduced hepatic expression was observed in genes important for oxidative phosphorylation and fatty acid metabolism (Fig. 2B, Supplementary Table 2), as well as genes required for drug metabolism, steroid hormone biosynthesis, and degradation of certain amino acids (Fig 2A, Supplementary Table 2), generally corresponding to genes found in clusters 4 and 5 in Fig. S3A. Reductions in hepatic expression of these pathways were primarily restricted to 6 and 10 dpi, and expression of these genes returned to baseline levels from 17 to 36 dpi. Suppression of hepatic oxidative phosphorylation and fatty acid metabolism during *C. rodentium* infection was previously linked to an infection-induced increase in bile metabolites, including itaconate, which has antimicrobial and immunomodulatory activities[16]. We propose that the localization of metabolic suppression to the liver, rather than the colon, reflects a facet of the systemic response to enteric infection.

**Fig. 2.**
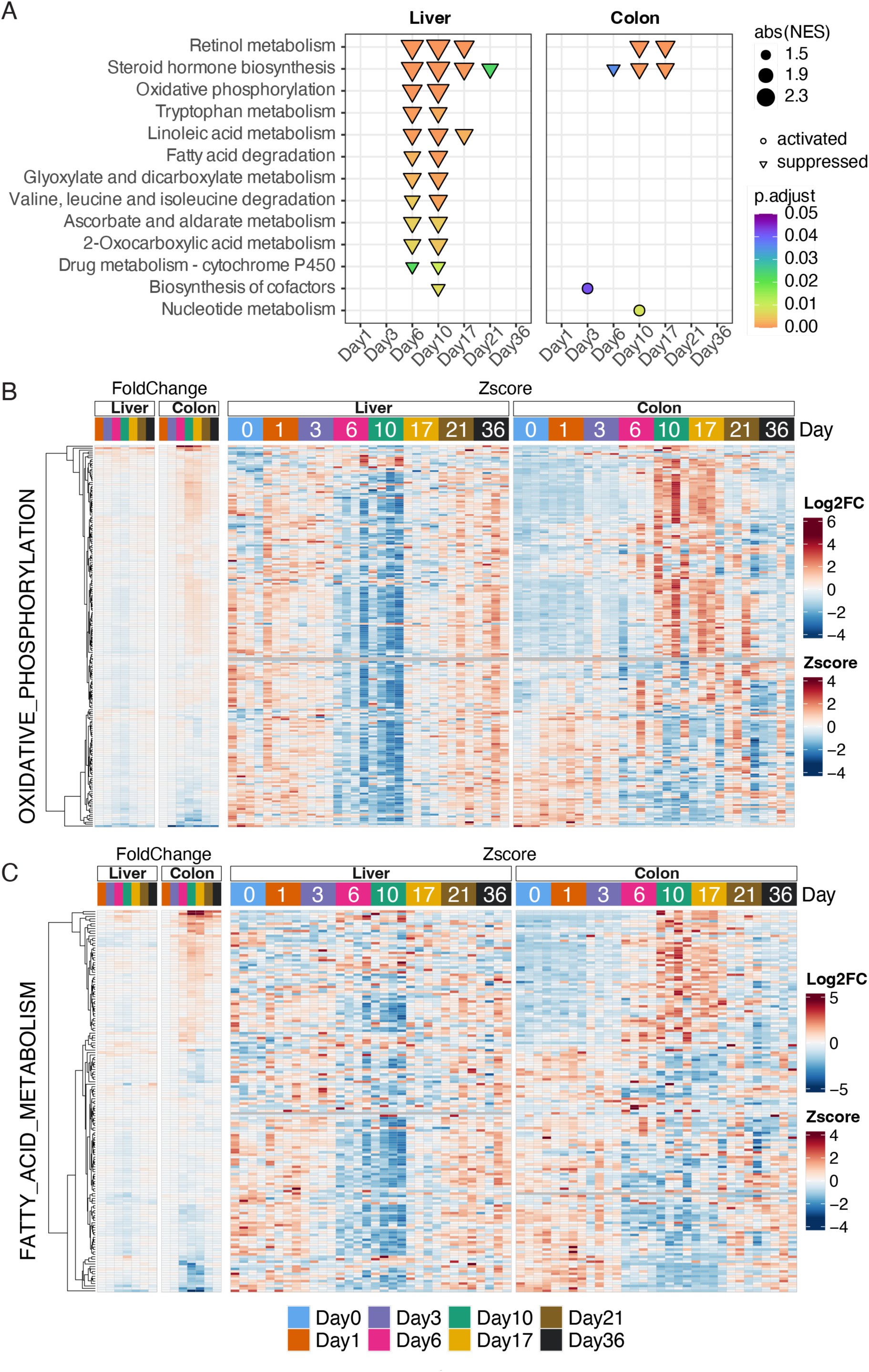
Expression of hepatic metabolic pathways is down-regulated during C. rodentium infection. **A.** Dot plots representing activated (circle) or suppressed (triangle) pathways in the liver and colon during C*. rodentium* infection. Dot size and color scale indicate the normalized enrichment score (NES) and adjusted *p*-value, respectively. KEGG Pathways that meet cutoff with adjusted p-value <0.01 in either the liver or colon epithelium at least one time point are shown. Complete pathway analysis data are included in Supplementary Table 2. **BC.** Heatmaps showing average log_2_ fold change relative to day 0 sample (left) and z-score (right) of genes in OXIDATIVE_PHOSPHORYLATION (B) and FATTY_ACID_METABOLISM (C) pathways. The color code on top represents the time points (days) after infection.

### Enteric infection results in a long-lasting change in colonic gene expression linked to persistent immune cell infiltration

GSEA using MSigDB [34] also revealed activation of pathways linked to cell death (apoptosis) and proliferation (e.g., E2F targets, G2M checkpoints, and mitotic spindle) in the liver and colonic epithelium, concurrent with infection and resolving around day 17 (Fig. 3A, Supplementary Table 2), consistent with previous observations that *C. rodentium* infection is associated with tissue damage in both the colon and liver[28]. Furthermore, potent activation was detected in genes related to innate immune signaling, including type I and II interferon signaling, IL6/Jak/Stat signaling, and TNFα signaling (Fig 3AB, Supplementary Table 2). In general, activation of these pathways was more consistent and long-lived in the colonic epithelium than in the liver. While heightened expression of these pathways was no longer apparent in the liver after 10 dpi, elevated expression of the genes for many of these pathways persisted until at least 36 dpi in the colon, suggesting that some of these genes/pathways represent the long-lived response to infection that was evident in the PCA analysis (Fig 1 FGH).

**Fig. 3.**
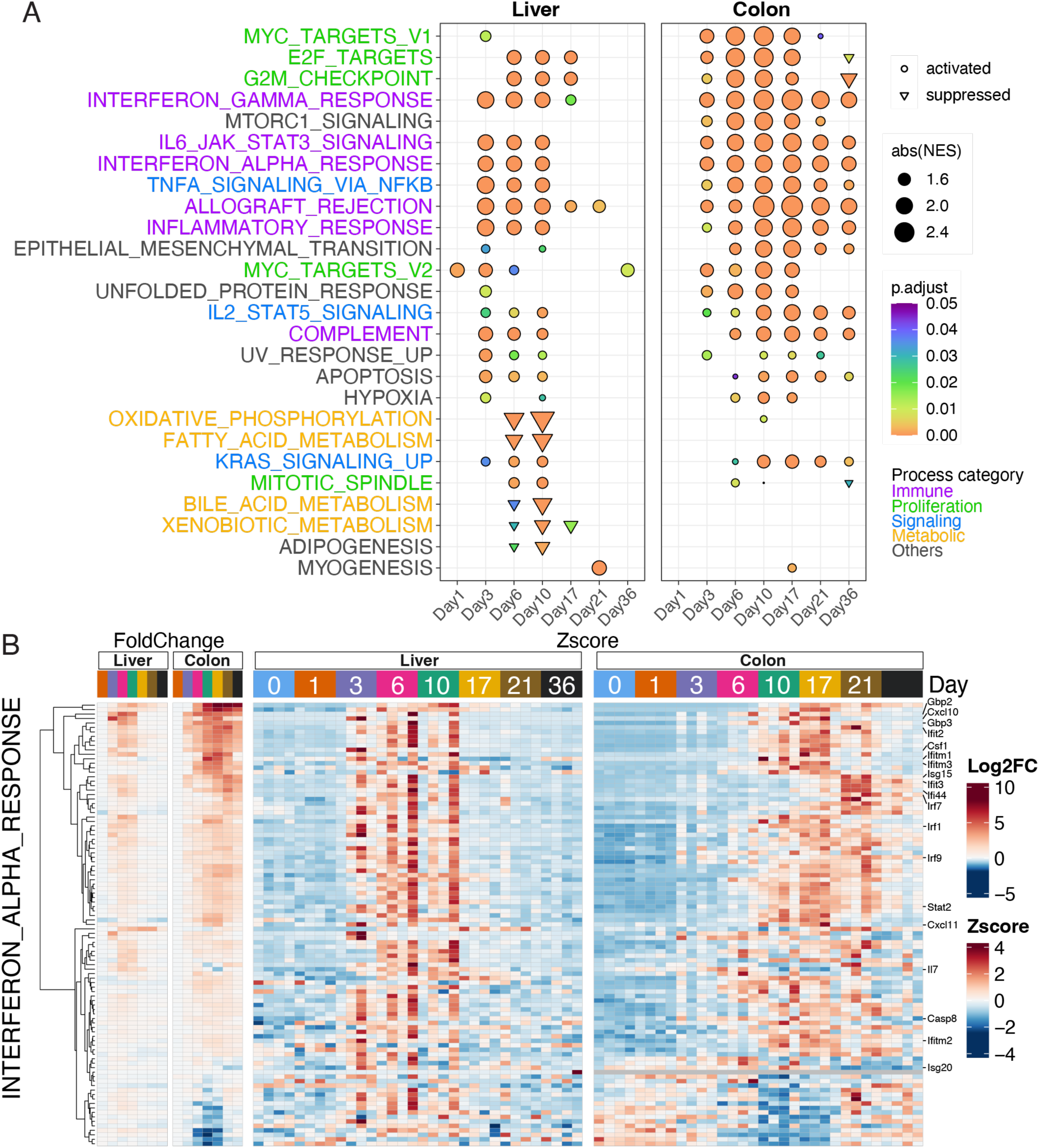
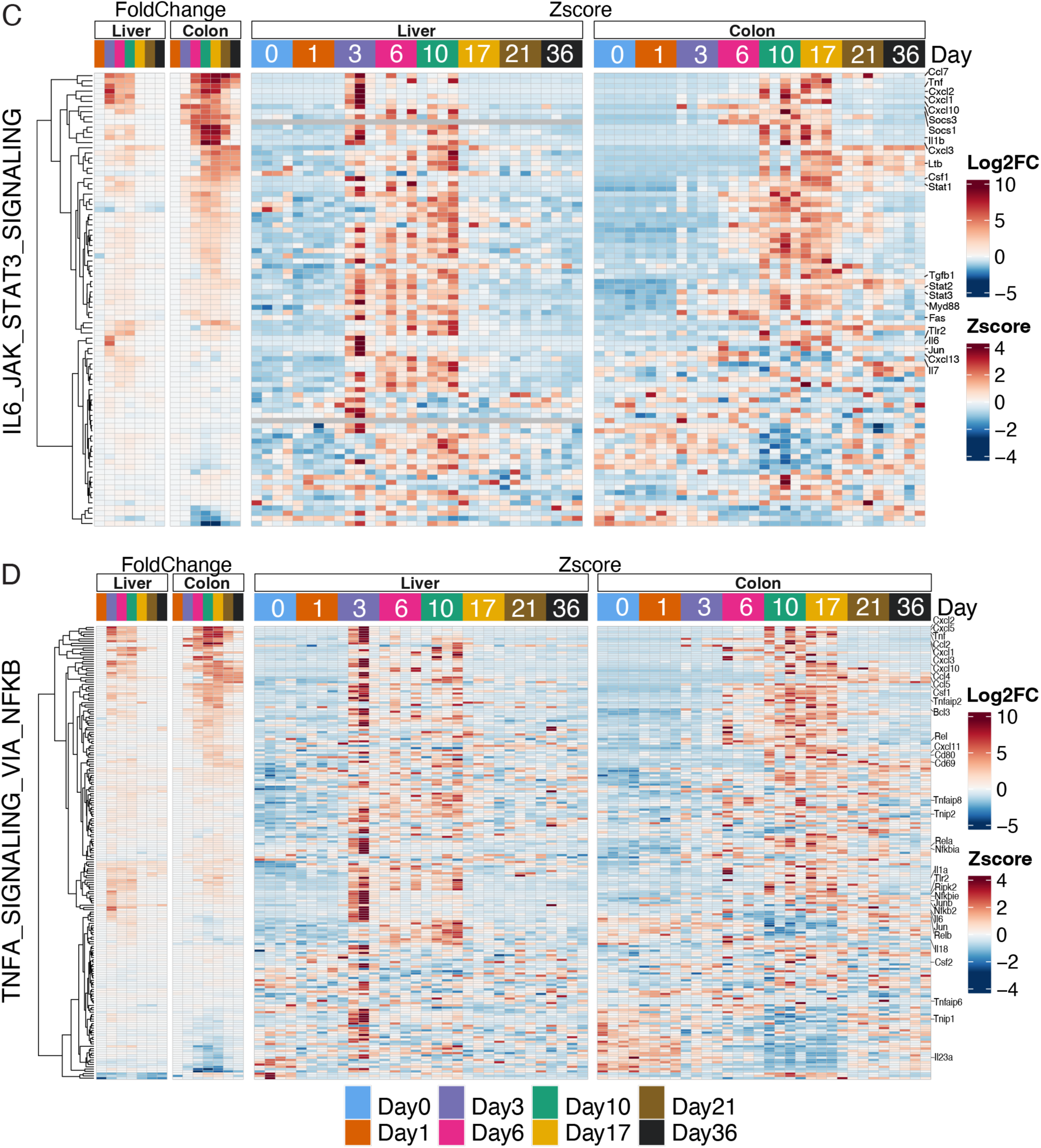
Temporal patterns of activation of immune signaling pathways in the liver and colon epithelium in response to C. rodentium infection. **A.** Dot plots representing activated (circle) or suppressed (triangle) pathways in the liver and colon by *C. rodentium* infection. Dot size and color scale indicate the normalized enrichment score (NES) and adjusted *p*-value, respectively. MSigDB Pathways that meet cutoff with adjusted p-value <0.01 in either the liver or colon epithelium at least one time point are shown. Pathway names are colored in the Processed category listed in Liberzon et al.[34] Complete pathway analysis data are included in Supplementary Table 2. **B-D.** Heatmaps showing average log_2_ fold change relative to day 0 sample (left) and z-score (right) of genes in INTERFERON_ALPHA_RESPONSES (B), IL6_JAK_STAT_SIGNALING (C), and TNFA_SIGNALING_VIA_NFKB (D) pathways. The color scale for the “FoldChange” plot represents the mean log_2_ fold change within each sample group, whereas the color scale for the “Zscore” plot shows the gene expression z-score for individual tissue samples. Each row of the heatmap depicts the expression of the same gene across time points in the two tissues. Selected gene names are labeled.

Multiparameter flow cytometry was used to investigate the hypothesis that infection-stimulated immune cell infiltration contributes to the increased expression of immune signaling pathways in the colon and liver (Fig. S4A-F). In the liver, infiltration of neutrophils (CD11b+, Ly6c+, Ly6g+), monocytes (CD11b+, Ly6c+, Ly6g-), and monocyte-derived macrophages (CD11b+, Ly6c+, Ly6g-, CD64+, F4/80+) peaked ∼10 dpi and returned to baseline 17 to 30 dpi (Fig. 4 AB, Fig. S4A). Invasion of these innate immune cells was more pronounced in the colon, both in the lamina propria (LP; Fig 4. DE, Fig. S4C) and epithelium (IEL; Fig. 4 GH, Fig. S4E). In addition to the larger magnitude of increase in the abundance of these cells in the colon compared to the liver, the peak abundance of these cells also occurred later in the colon (17 dpi) compared to the liver (10 dpi). The relatively delayed peak in the influx of innate immune cells in the colon, which occurred after the peak in pathogen burden, may reflect ongoing residual pathogen clearance and tissue repair in response to pathogen-related damage[35,36] (Fig. 4 BEH). In the colon, there was also a marked increase in the abundance of CD4 T cells (>30-fold) and CD8 T cells (>10-fold), which also peaked at 17 dpi and persisted until at least 30 dpi (Fig. 4 DFGI). The T cells infiltrating the IEL were primarily CD28+ CD44+ memory T cells (Figure S4GH), potentially accounting for the elevation of gamma interferon signaling in the colon[37], which persisted even after pathogen clearance (Fig. 1A). Thus, *C. rodentium* infection results in long-lived residence of memory CD4 and CD8 T cells in the colon, which could augment protection against future infection[38].

**Fig. 4.**
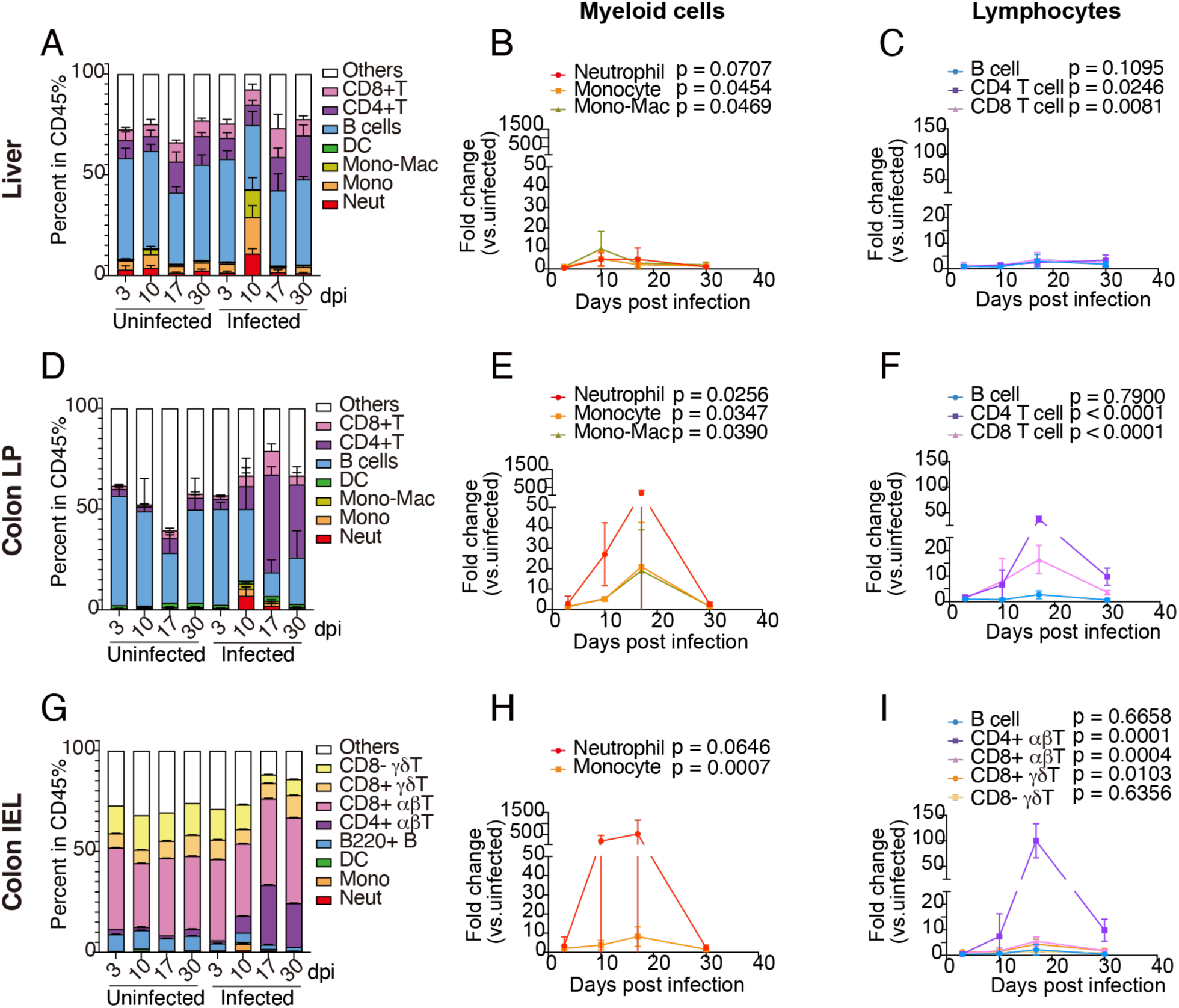
Temporal course of immune cell profiles in the liver, colonic lamina propria, and colonic epithelium in response to C. rodentium infection. **A-I**. Changes in immune cell populations in the liver (A-C), colon lamina propria (LP, D-F), and colonic epithelial tissue (IEL, G-I). **A. D. G.** The fraction of CD45+ immune cells in the liver (A), colonic LP (D), and colonic IEL (G). X-axis and Y-axis indicate the days post inoculation (Days) in uninfected or infected group, and percentage of each cell type among CD45+ cells, respectively. Color code shows cell types. **B. E. H.** Fold changes of the number of neutrophils, monocytes, and monocyte-derived macrophages (Mono-Mac) compared to the time-matched uninfected control in the liver (B), colonic LP (E) and colonic IEL (H). Color code represents cell types. Two-way ANOVA; compared to uninfected control. **C. F. I.** Fold changes of the number of indicated lymphocytes compared to the time-matched uninfected control in the liver (C), colonic LP (F) and colonic IEL (I). Color code represents cell types. Two-way ANOVA; compared to uninfected control. mean ± SD. n = 5/group.

### Enteric infection induces an early liver gene signature linked to a self-limited systemic inflammatory state

Temporal analysis captured an early hepatic response to enteric infection 3 dpi (Fig. 1E, PC2) that occurred prior to infection-related immune cell infiltration into the liver (Fig. 4 ABC). Unlike the prolonged immune response in the colon (Fig. 3AB), this early hepatic response was self-limited, reversing 6 dpi, before the peak of enteric colonization and detection of *C. rodentium* in the liver 6-10 dpi (Fig. 1A), and prior to the resolution of the liver apoptosis signal 17 dpi (Fig. 3A). Analysis of pathways captured by liver PC2 suggests that this early, self-limited wave of gene expression may modify the systemic immune state. At 3 dpi, the top 100 genes by loading-score in liver PC2 were nearly all involved in immune signaling, including genes related to TNF, IL17, NFκB, and other cytokine signaling (Fig 5A).

**Fig. 5.**
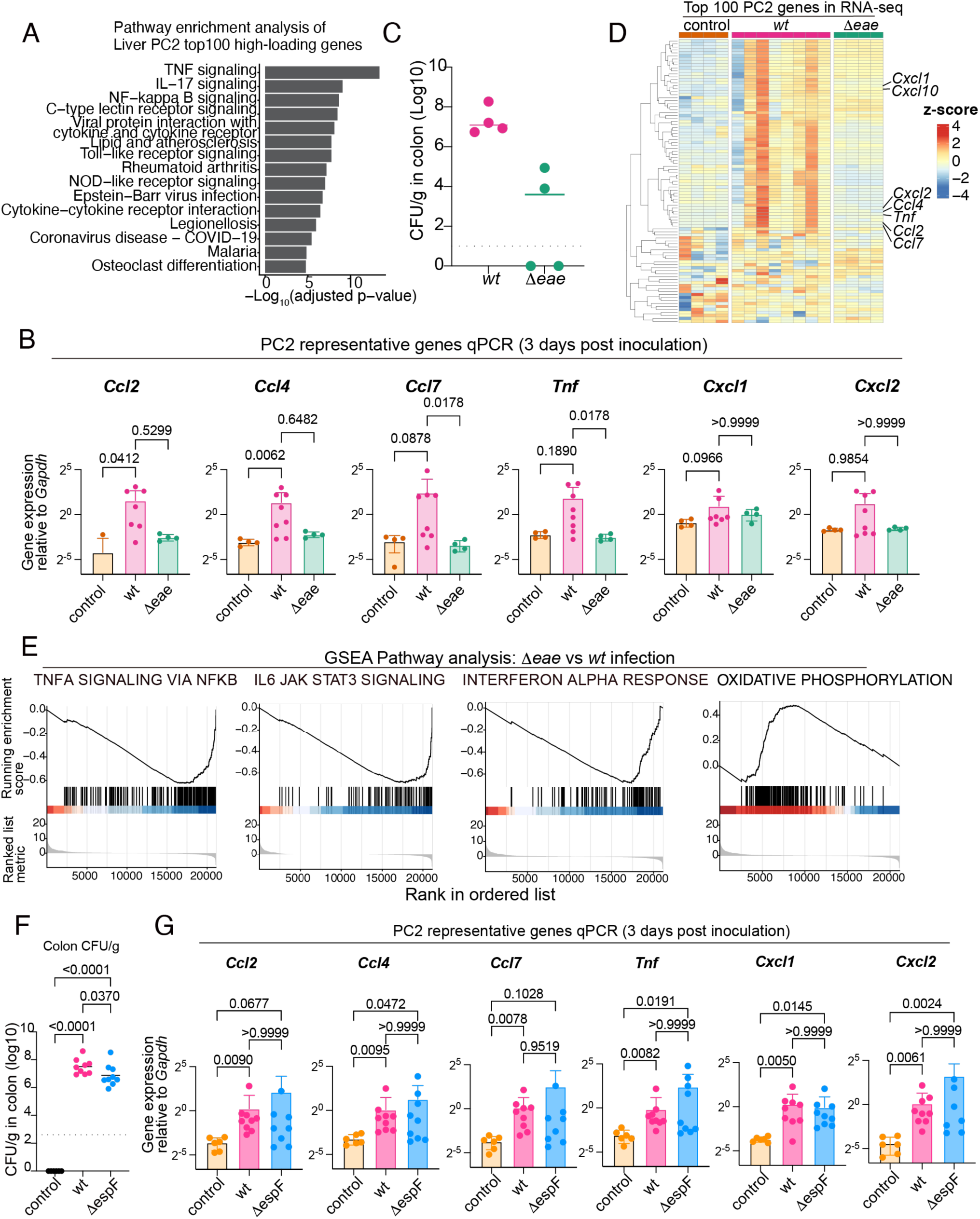
Early hepatic immune signaling in response to C. rodentium infection is intimin dependent. **A.** Pathway enrichment analysis of the top 100 genes with high PC2 loading score 3 days post infection with wildtype *C. rodentium*. The x-axis shows -log_10_ transformed adjusted p-value. Top 15 pathways are shown. **B. G.** qPCR analysis of representative PC2 gene expression in the liver in animals infected with wild type (wt), Δ*eae* or Δ*espF C. rodentium* 3 days post inoculation. Y-axis shows the relative value to *Gapdh* expression. **B.** Control (n = 4), wt-infected (n = 8), and Δ*eae*-infected (n = 4). **G.** Control (n = 6), wt-infected (n = 9), and Δ*espF*-infected (n = 9). Kruskal-Wallis test with Dunn’s multiple comparisons test. **C.** Wt or Δ*eae C. rodentium* burden (CFU/g) in the colon on 3 days post inoculation. Control (n = 4), wt-infected (n = 8), and Δ*eae*-infected (n = 4). **D.** Liver RNA-expression of top 100 PC2 genes in uninfected control, wt-infected (wt), and Δ*eae*-infected (Δ*eae*) animals. Z-score of RNA expression is shown in the color scale. Each column represents an individual animal sample. Uninfected control datasets and four wt-infected samples originate form the time-series transcriptome dataset in Fig. 1. Z-score conversion was performed after data integration, batch correction, and normalization, as described in the Materials and methods. **E.** Gene set enrichment analysis (GSEA) for liver RNA expression comparing Δ*eae* vs. wt *C. rodentium*-infected animals. Experiments for Fig. 5B and Fig. 5CD used different batch of animals. Data from uninfected control and four wt-infected animals originate from data on 3 dpi in time course experiment (Fig. 1). **F.** Wt or Δ*espF C. rodentium* burden (CFU/g) in the colon on 3 days post inoculation. One-way ANOVA after lognormal conversion with Holm-Šídák’s multiple comparisons test. Dotted line shows the limit of detection. Control (n = 6), wt-infected (n = 9), and Δ*espF*-infected (n = 9)

The magnitude of the early hepatic response described by PC2 was heterogeneous across animals (Fig 1E). Given that this response was observed only at 3 dpi, we propose that it occurs within a limited time window that was not fully captured in all mice. Quantitative PCR of PC2 marker genes (*Ccl2*, *Ccl4*, *Ccl7*, *Tnf*, *Cxcl1*, and *Cxcl2*) in additional animals confirmed a similar heterogeneous pattern of activation 3 dpi (Fig. 5B), with increased transcripts detected in 4 out of 8 infected animals compared to uninfected controls.

Mice were challenged with a *C. rodentium* mutant (Δ*eae*) unable to attach to the host epithelium[39] to differentiate whether the early hepatic PC2 response is dependent on the pathogen’s virulence program, which primarily acts downstream of intimin-mediated attachment[20], or by the initial bolus of bacteria and microbe-associated molecules from the intragastric gavage. Consistent with previous report[39], *C. rodentium* lacking the gene encoding the adhesin intimin (Eae) failed to expand in the intestine (Fig. 5C). Furthermore, inoculation with the Δ*eae C. rodentium* mutant did not cause the same early hepatic response 3 dpi as wild-type infection (Fig 5. BDE), including fewer transcripts in the liver from PC2 marker genes detected by quantitative PCR (Fig. 5B) and RNA-seq (Fig. 5D), as well as PC1 genes (Fig. S5). Comparison of liver transcripts 3 dpi following Δ*eae* or wild-type *C. rodentium* challenge by GSEA leading-edge analysis further highlights that activation of inflammatory pathways, including TNFα and IL6 signaling, requires enteric pathogenesis, i.e., the intimin-mediated pathogen-host interactions. Thus, we conclude that the early hepatic PC2 response is triggered by *eae*-dependent enteric infection and cannot be explained solely by the bacterial bolus in the inoculum.

*C. rodentium* infection increases epithelial permeability by disrupting tight junctions through the activity of type III effectors, including EspF and Map[22,23]. *C. rodentium* strains lacking EspF (Δ*espF*) have been reported to colonize the colon as efficiently as wild-type bacteria with reduced disruption of tight junctions[22]. Challenge with an Δ*espF* strain was used to investigate whether tight junction disruption is required for the early hepatic PC2 response. There was a mild colonization defect in the Δ*espF* strain (4.3-fold) (Fig. 5F), and dissemination to the liver was largely not observed in both the wild-type and Δ*espF* strains at 3 dpi (Fig. S6). Quantitative PCR analysis at 3 dpi demonstrated that infection with Δ*espF C. rodentium* increased PC2 marker gene transcripts in the liver as high as infection with the wild-type strain (Fig. 5G), suggesting that disruption of the tight junction does not account for this liver response.

To determine the consequences of early hepatic PC2 activation on systemic immune tone, we measured serum concentrations of 46 cytokines in samples from the same animals used in the gene expression time-course experiments. These matched data revealed that the timing of liver gene expression strongly predicts the systemic abundance of a set of corresponding cytokines/chemokines (Fig. 6 ABC, red), including a set of ‘early hepatic response genes’ that peaked in liver expression and corresponding serum protein abundance at 3 dpi, and returned towards baseline thereafter. In contrast, only the expression of a single transcript (TIMP-1) in the colonic epithelium significantly predicted systemic protein abundance (Fig. 6D). The set of early hepatic response genes included TNF-α and IL-6, key mediators of acute-phase protein expression, as well as IP-10 (Cxcl10), KC (Cxcl1), MCP-1 (Ccl2), and MIP-1β (Ccl4). The magnitude of expression of these early hepatic response genes decreased in the liver from 6-36 dpi, whereas expression of the same set of immune signaling genes increased in the colonic epithelium and peaked 10-17dpi (Fig. 6B). However, this localized burst of colonic immune gene expression was not reflected in systemic concentrations of the early hepatic response factors, suggesting that the liver coordinates the systemic concentration of these proteins in response to enteric infection.

**Fig. 6.**
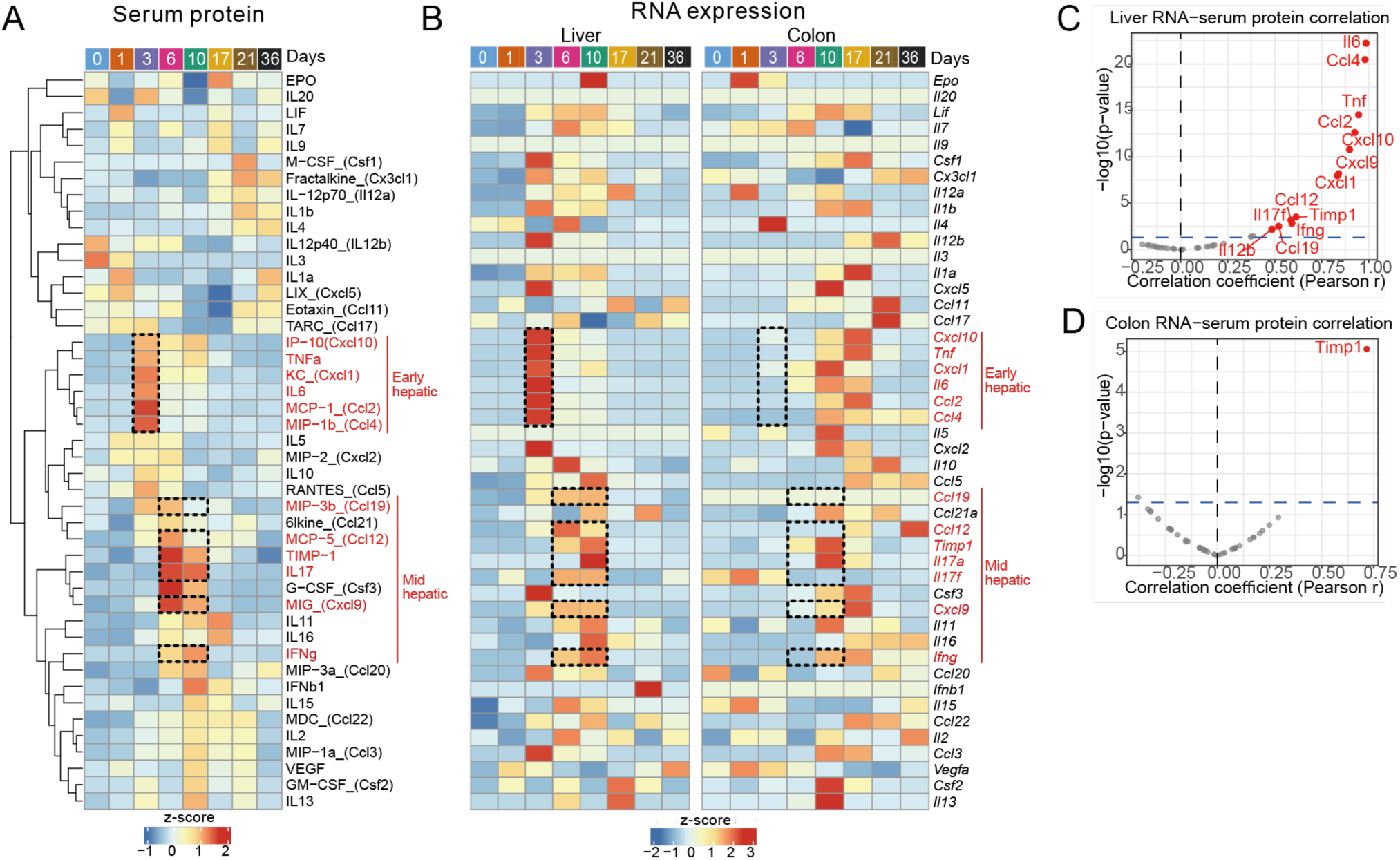
Temporal correlation between abundance of hepatic cytokine RNAs and serum cytokine protein abundance. A. Protein abundance of cytokine and chemokine in serum over the course of infection. The mean z-score of the abundance of each protein is shown in color. The color code at the top indicates time points (days) after infection. n = 5/ Days. B. RNA expression of cytokine and chemokine in the liver and colon. The mean z-score of RNA expression is shown in color. The color code at the top indicates time points (days) after infection. n = 4/Days. **C,D.** Correlation between the abundance of serum protein and RNA expression in the liver (C) or colonic epithelium (D). X-axis and y-axis indicate Pearson’s correlation coefficient and -log10 scaled p-value, respectively. Genes with *p* < 0.01 are shown in red. n = 32.

Cytokine measurements also revealed increased abundance of several serum proteins at 6-10 dpi, which correlated with corresponding increases in hepatic gene expression (Fig. 6A-C). These ‘mid hepatic response’ genes included MIP-3β (Ccl19), MCP-5 (Ccl12), TIMP-1, IL17, MIG (Cxcl9), and IFNγ. Elevated expression of these genes was also observed in the colon (Fig. 6B); however, expression in the colonic epithelium was not correlated with systemic cytokine abundance, except for TIMP-1 (Fig. 6D). Together, these observations provide correlative evidence that waves of hepatic gene expression control the systemic response to intestinal infection, including limiting the acute-phase protein response to an early stage of infection.

## Discussion

Here, to uncover mechanisms that account for systemic responses to enteric infection, we compiled matched timeseries data that included *C. rodentium* intestinal burden along with gene expression profiles in the liver and colonic epithelium. Pairing the kinetics of pathogen burden with those of organ-specific gene expression profiles revealed several unexpected insights into host responses to intestinal infection, including likely hepatic drivers of systemic responses. First, there were more diverse ‘early’ changes in hepatic vs. colonic epithelium gene expression profiles during the first three days of infection, prior to peak pathogen burden and disease in the colon. These early changes primarily depended on the pathogen’s virulence program and were not observed in mice inoculated with a *C. rodentium* Δ*eae* mutant. Second, the temporal patterns of RNA abundance in the liver and colonic epithelium accompanying enteric infection were largely distinct. Finally, the timing of hepatic changes in transcripts linked to cytokines and immunomodulatory factors closely mirrored changes in serum cytokine abundance. Particularly noteworthy was the transient elevation of transcript levels and serum abundance of IL-6 and TNFα at 3 dpi; these cytokines control the expression of acute-phase response proteins[25,40]. Together, our data suggest that early and self-limited hepatic responses to enteric infection control a key aspect of the kinetics of systemic immune tone during the course of gut-localized infection. Furthermore, our findings demonstrate the power of matched timeseries data for deciphering the multi-compartment connections between localized infections and systemic responses.

Monitoring hepatic gene expression profiles throughout the entire course of *C. rodentium* murine colitis revealed a transient signal that peaked 3 dpi and largely returned to baseline by 6 dpi (Fig 1, PC2). This pattern preceded and resolved before the peak of pathogen burden; *C. rodentium* was not detected in the liver 1-3 dpi, while colonic burden peaked 6-10dpi. This early hepatic signal was strongly weighted toward genes related to innate immune responses, particularly cytokine signaling, such as TNFα (Fig 5A). Although liver pathogen burden cannot account for the burst of gene expression, the increase in immune signaling was relatively muted when an intimin mutant was used to inoculate mice (Fig 5D), indicating that the *C. rodentium* intimin-dependent virulence program is critical for triggering hepatic immune signaling. In contrast, deletion of the gene *espF*, which is required to disrupt tight junctions (PMID: 16548889), did not affect the early hepatic transcriptomic response (Fig. 5G). Thus, signals independent of tight junction disruption, including those from tissue-resident immune cells and translocation of disruption-independent microbial-associated patterns, likely account for the early hepatic response.

The signal linking *C. rodentium* infection to activation of the early hepatic immune response remains an important area for further investigation, but previous work has established that *C. rodentium* intimin-dependent interactions with the epithelium can stimulate proinflammatory cytokine production[20,41], myeloid and T cell recruitment[42], and mucosal adaptive immune responses[41,43]. Thus, it’s possible that host-derived molecules produced in the colon in response to infection and delivered to the liver via the portal vein activate early hepatic immune signaling. Given that this response occurs at 3 dpi, prior to substantial infection-dependent immune cell infiltration into the colon (Fig. 4 D-I), it is possible that the signal originated from resident intestinal immune cells in the lamina propria and not captured in the transcriptomic profiling of the colonic epithelium conducted in this study. Furthermore, given the tissue damage caused by prolonged IL-6 signaling [2], interrogating the mechanisms that limit early hepatic immune activation to a short window could illuminate the homeostatic mechanisms that prevent prolonged inflammation and damage after enteric infection.

Regardless of the mechanisms underlying the self-limited early hepatic immune signaling, this process likely shapes the kinetics of the acute phase response during gut-localized infection. Here, in response to enteric infection, we observed a single spike in hepatic IL-6 and TNFα expression and a concomitant increase in their corresponding serum cytokine levels at 3 dpi. Exogenous IL-6 or TNFα is sufficient to trigger an early wave of production of acute phase proteins [24,26]. Thus, we propose that the kinetics of hepatic cytokine expression underlie the temporal course of the early systemic response to enteric infection. Dysregulation of the yet to be defined pathways that underlie the self-limited expression of these cytokines could lead to chronic inflammatory sequelae of intestinal infection, such as Reiter’s disease [44].

Several aspects of the trajectories of the changes in gene expression profiles detected in the colonic epithelium differed from those observed in the liver. Although the colon is the chief site of *C. rodentium* colonization, gene and protein expression changes there [45,46], which were ultimately more diverse compared with the liver, were delayed. Moreover, changes in colonic epithelial gene expression profiles endured beyond the period of maximal pathogen burden 10 dpi. For example, the number of colonic DEGs remained similar on 10 and 17 dpi, even though at the later time point some animals had already cleared the pathogen. Moreover, even at 36 dpi, colonic gene expression profiles remained noticeably different from their baseline, for example, long-lasting activation of interferon gamma responses (Fig. 3A), consistent with prior proteomic studies [45]. Presumably, the persistent changes in expression profiles observed in the colon vs the liver reflect the more pronounced tissue damage or responses to bacteria-derived products in the colon, which results in infiltration of immune cells that mediate pathogen clearance, tissue repair, and immune memory of infection [37,38,47–51].

Besides differences in the kinetics of hepatic and colonic responses to enteric infection, the identities of the responding genes and the systemic consequences of these responses appear to differ. For example, in Figure 3B, almost the entire set of genes in the MSigDB interferon alpha response pathway in the liver are activated 3-10 dpi, whereas in the colon, there is less comprehensive activation of the genes in this pathway. These observations suggest that distinct regulatory processes mediate inflammatory responses to infection in these two tissues. Also, while marked increases in pathways linked to inflammation were observed in both the liver and colon, elevated expression of these pathways persisted longer in the colon and may reflect the influx of innate immune cells like neutrophils and monocytes, as well as memory CD4 and CD8 T cells in the colon (Fig 4). Although expression of many inflammation-linked genes in the liver and colon increased as early as 3 dpi, these signaling pathways remained elevated in the colon through 36 dpi, when serum cytokine levels had largely returned to baseline (Fig 6). Together, these observations suggest that the localized inflammatory response in the colonic epithelium is not as potent a modulator of systemic immune tone as the liver. Thus, although the initial intimin-dependent tissue damage most likely originates in the intestine, liver gene expression correlates strongly with systemic inflammation, suggesting that it functions as an amplifier and coordinator of systemic responses to enteric infection. The elevation of systemic inflammatory cytokines likely control intestinal infection by processes such as increasing myelopoiesis in the bone marrow[52], increasing vascular permeability for immune cell infiltration[53,54], elevation of body temperature[55], and even changes animal behavior[56].

There was also more pronounced reductions in liver versus colon expression of genes linked to metabolism, including cytochrome-linked drug metabolism, oxidative phosphorylation, and fatty acid metabolism at peak pathogen burden (6-10 dpi) (Fig 2). Similar reductions of metabolic gene transcripts in the liver were previously observed [16], but the systemic consequences of these changes have not been evaluated. Reduction in oxidative phosphorylation and fatty acid metabolism may enable the liver to devote more resources toward proliferation of immune cells and hepatocyte regeneration, since these processes depend on anabolic glycolysis [57–59]. Our transcriptional profiling of the colon showed less suppression of metabolism than previous proteomic analyses[45,46]. These discrepancies between proteomic and transcriptomic analyses suggest that the abundance of colonic epithelial proteins may be regulated post-transcriptionally.

Following hepatic gene expression throughout infection was valuable because it revealed that a discrete burst of gene expression at 3 dpi appears to coordinate critical systemic immune responses to enteric infection. This early response was not detected in all animals. For example, in Fig 1C,E, changes in PC2 3 dpi were observed in 3 of 4 animals. Such heterogeneity could be due to animal-to-animal variability in the magnitude of the PC2 response. Alternatively, and not mutually exclusive with this idea, we favor the hypothesis that there are animal-to-animal differences in the kinetics of inter-organ signaling. Thus, since the expression pattern captured by PC2 is temporally discrete, changes in expression of PC2-linked genes could occur in all animals, but be missed in some animals at the time of sampling.

Overall, our study highlights the value of matched timeseries data for deciphering interorgan crosstalk and its systemic consequences. The efficacy and damage caused by systemic immune responses depend on their timing, and these dynamics are often missed by single-timepoint analyses.

## Limitations

This study uses time series data from mice infected with an enteric pathogen to link an early transcriptional response in the liver to systemic cytokine responses. While this temporal analysis provides strong correlational evidence linking these events, this work does not provide direct experimental evidence connecting the events in the liver to protein levels in the blood. Further studies are required to identify the molecular messengers, including from the cecum, that trigger the early hepatic responses and to measure the impact of preventing its delivery to the liver on systemic immune tone.

## Materials and methods

### Mice

Specific-pathogen-free (SPF) C57BL/6 J female mice aged 8-10 weeks mice were purchased from Jackson Laboratory (strain #000664;Bar Harbor, ME, USA). Mice were co-housed and maintained on a 12-hour light/dark cycle with access to standard chow in the SPF animal facility at the Brigham and Women’s Hospital. All animal studies were conducted in a biosafety level 2 (BSL2) facility at the Brigham and Women’s Hospital. Experiments were performed in accordance with protocols reviewed and approved by the Brigham and Women’s Hospital Institutional Animal Care and Use Committee (protocol 2016N000416) and in compliance with the Guide for the Care and Use of Laboratory Animals.

### Bacterial Strains

A spontaneous streptomycin-resistant mutant of *C. rodentium* strain ICC168, previously known as *Citrobacter freundii* biotype 4280 (ATCC 51459), was used here [30]. *C. rodentium* was grown at 37 °C in LB broth with shaking or on solid LB agar. A Δ*eae* and Δ*espF* mutants of *C. rodentium* strain ICC168 were created by standard allele exchange methods as described previously [60]. Briefly, PCR-amplified fragments corresponding to regions upstream and downstream of the *eae* or *epsF* coding sequence were assembled using NEBuilder HiFi DNA Assembly Master Mix (NEB) to generate a deletion construct. This construct was introduced into *C. rodentium* via conjugation using *E. coli* strain MFDpir [61]. The right mutation and the lack of unintended mutations were checked by comparing to the genome sequence of the wild type strain using the whole genome sequencing service provided by Plasmidsaurus (South San Francisco, CA).

### Mouse infection

*C. rodentium* was prepared for oral inoculation by resuspending frozen cells in liquid LB broth and expanding the culture overnight at 37 °C with shaking. Then, bacteria were pelleted, resuspended in PBS, and kept at room temperature until inoculation. Mice were deprived of food for 6 hour prior to inoculation, and then lightly anesthetized with isoflurane inhalation and orally inoculated with 5×10^9^ CFU of *C. rodentium* in 100 ul PBS using a 18G flexible feeding needle (Braintree Scientific). Following inoculation, the doses were corroborated by serial dilution and plating. The animals’ clinical condition (e.g weight, activity level, and fur appearance) and fecal appearance were monitored during infection. For quantifying *C. rodentium* burden in fecal pellets or organs, samples were homogenized in PBS using a bead beater (BioSpec Products, Inc) with 2 stainless-steel 3.2 mm beads. Homogenized samples were plated on LB plates containing 200 μg/mL streptomycin and incubated at 37 °C overnight prior to counting CFU.

### RNA preparation from colonic epithelial cells and liver

Colons were collected, opened longitudinally, and washed with 1 mL of Gut wash media (RPMI1640 supplemented with 2% FBS, 10 mM HEPES, 100 μg/mL Penicillin-Streptomycin and 50 μg/mL gentamicin) in 15 mL tubes. Tissues were then rinsed in ice-cold HBSS and cut into ∼2 cm pieces. The tissue pieces were incubated in 10 mL of Epithelial dissociation solution (HBSS containing 10 mM EDTA, 2% FBS, 10 mM HEPES, and 100 μg/mL Penicillin-Streptomycin) in 50 mL tubes at 37°C for 5 min on a shaker at 125 rpm. Tubes were then transferred to ice and incubated for 5 min, gently shaken 10 times, and briefly vortexed for 2 seconds. The tissue pieces were transferred to 10 mL fresh Epithelial dissociation solution in new 50 mL tubes and incubated again at 37°C for 25 min on a shaker at 125 rpm. Tubes were then placed on ice for 5 min, gently shaken 15 times, and vortexed for 10 seconds. Tissue pieces were removed using forceps, and the dissociated cell suspension was transferred to fresh 15 mL tubes and centrifuged at 300 xg for 5 min. The supernatant was discarded, and cell pellets were resuspended in 2 mL of TRIzol (Thermo Fisher). FACS analysis suggested that the majority of the cells in this epithelial fraction were Epcam-positive (epithelial cells), but some intraepithelial immune cells were also detected. In principle, there could be contamination from immune cells and stromal cells derived from the mucosal lamina propria. RNA was extracted according to the manufacturer’s instructions for TRIzol. The liver was collected and homogenized in 1ml of TRIzol, and RNA was extracted according to the manufacturer’s instructions. The quality of all RNA samples was assessed using an Agilent TapeStation. The RNA integrity number (RIN) were above 8.0 for the liver samples, except for Liver-day10-1, which had a RIN of 7.3, and above 9.0 for the colon epithelium samples.

### RNA-seq and data analysis

RNA-seq libraries were prepared using the NEBNext Ultra II Directional RNA Library Prep Kit (NEB) with NEBNext Poly(A) mRNA Magnetic Isolation Module (NEB). Libraries were sequenced on a NextSeq 550 with single-end runs of 75 bp, NovaSeq 6000 with single-end runs of 100 bp, or NextSeq2000 with paired-end runs of 2 × 75 bp or 2 x 151 bp. The average sequencing depth was approximately 40 million reads per sample, with a minimum of 20 million reads. Low-quality bases and the adaptors were trimmed using Trim Galore (v0.6.6) with a paired option. The reads were mapped to the mouse reference genome (mm10) using STAR v2.7.3a with default parameters. More than 85% of reads were uniquely mapped, as assessed based on the STAR log reports. The gene-level raw read count matrix was obtained using the featureCounts function in the subread (v2.0.0) using mouse GENCODE annotation M25 (GRCm38.p6). Differentially expressed genes were identified using the R-package DESeq2 (v1.36.0) with an adjusted *p*-value < 0.05 and an absolute log_2_ fold change of 1 relative to uninfected time 0 control animals, and the clustering was performed using the pam function in the R-package cluster (v2.1.4) with a parameter ‘k = 6’. Pathway analysis was performed by the R-package clusterProfiler (v4.4.4) with gene sets from msigdbr R-package (v7.5.1) or KEGG metabolism pathways. Database and version information for the pathway analysis have been added to the top row of the Supplementary Table S2. For the analysis and visualization of liver RNA-seq data in infection with Δ*eae* strain (Figure 5D), count table of day 3 data from time series analysis was combined with count table from wt or eae-infected animals. To compare liver transcriptome data on 3 dpi in time series data with those of animals infected with Δ*eae* mutant (Fig. 5D), batch correction was performed using Combat-seq[62] command in sva package (v3.35.2) with default settings. Data is visualized with pheatmap package (v1.0.13).

### Immune cell isolation from the colon and liver

Following collection of the colon, the tissue was opened longitudinally, and washed with 3 mL of Gut wash media in a 12-well plate. Tissues were then rinsed in ice-cold HBSS and cut into ∼2 cm pieces. The tissue pieces were incubated in DTT solution (HBSS containing 1mM DTT, 10mM HEPES, and 5% FBS) and filtered through a 70µm strainer. The tissue was then incubated in 10 mL of Epithelial dissociation solution (HBSS containing 15 mM EDTA, 10 mM HEPES, 2% FBS and 100 ug/mL Penicillin-Streptomycin) in 50 mL tubes at 37°C for 15 min on a shaker at 125 rpm. Tubes were briefly vortexed for 10 seconds. The aqueous phase was collected and resuspended with 44% Percoll and IEL was separated by adding 67% Percoll. The remaining tissue was then incubated in Enzyme mix buffer (RPMI1640 glutamax containing 10mM HEPES, 0.5% Collagenase D, 0.5% Dispase II, and 0.05% DNase I) at 37℃ for 1h. The digested tissue were then resuspended with 40% Percoll and LP was separated by adding 80% Percoll. The liver was squeezed through 1mm filter (pluriSelect 43-51000-01) and incubated in Enzyme mix buffer (RPMI1640 glutamax containing 10mM HEPES, 0.5% CollagenaseD, 0.5% DospaseII, and 0.05% DNase I). The digested tissue was filtered through a 70 µm filter, and then the red blood cells ware lysed by RBC lysis buffer (Roche, 11814389001). Remaining tissue pieces were resuspended with 36% Percoll and hepatic immune cells were separated by adding 72% Percoll. Separated IEL,LP, and hepatic immune cells were kept in FACS buffer (PBS containing 2 mM EDTA and 2% FBS) on ice till further analysis.

### Spectral Flow Cytometry

All the antibodies used in this study are listed in Table 1. Single-cell suspensions were blocked with an antibody against CD16/32 (2.4G2, Becton Dickinson) and then stained with either LIVE/DEAD fixable aqua cell stain kit (L34957, Invitrogen) or LIVE/DEAD Fixable Near-IR Dead Cell Stain Kit (L10119, ThermoFisher) and target antibodies. After incubation at room temperature for 30 min, cells were washed in PBS, fixed in 4% paraformaldehyde PBS for 10 min, washed in PBS, and analyzed with a Cytek Northern Lights spectral flow cytometer and analyses were performed with FlowJo software (v10.8).

**Table 1.** Antibodies for spectral flow cytometry analysis.

| Antibodies | clone | Vender | Cat. No. |
| --- | --- | --- | --- |
| Anti-CD45 Alexa Flour 700 | 30-F11 | BioLegend | 103128 |
| Anti-B220 PE-Fire 810 | RA3-6B2 | BioLegend | 103287 |
| Anti-CD3e Super Bright 600 | 145-2C11 | Thermo Fisher Scientific | 63-0031-82 |
| Anti-F4/80 PE-Fire 640 | QA17A29 | BioLegend | 157320 |
| Anti-CD11c PE-Cy7 | HL3 | BD Bioscience | 558079 |
| Anti-MHC II (IA-IE) Alexa Flour 488 | M5/144.15.2 | BD Bioscience | 562352 |
| Anti-CD11b APC | M1/70 | BD Bioscience | 553312 |
| Anti-Ly6C PerCP-Cy5.5 | HK1.4 | BioLegend | 128012 |
| Anti-Ly6G APC-Cy7 | 1A8 | BioLegend | 127623 |
| Anti-CD64 PE | X54-5/7.1 | BD Bioscience | 558455 |
| Anti-TCRβ PE-CF 594 | H57-597 | BD Bioscience | 562841 |
| Anti-TCR$\gamma/\delta$ APC | GL3 | BioLegend | 118143 |
| Anti-CD4 Super Bright 702 | RM4-5 | Cytex | 67-0041-80 |
| Anti-CD8a Super Bright 780 | 53-6.7 | Cytex | 78-0088-42 |
| Anti-CD28 PE-Cy7 | E18 | BioLegend | 122014 |
| Anti-CD44 Brilliant Violet 650 | IM7 | BioLegend | 103049 |

### Serum cytokine analysis

Blood samples were collected from either the tail vein or the inferior vena cava. Blood was collected into Eppendorf tube and was left at room temperature until it coagulates. Samples were subsequently centrifuged at 845 xg for 15 min to separate the serum. Serum cytokine levels were quantified using a multiplex immunoassay platform by Eve Technologies (Calgary, Canada). Data were analyzed using proprietary software provided by Eve Technologies, and cytokine concentrations were reported in pg/mL.

Pearson’s correlation coefficient was calculated as follows; RNA count table was converted into transcript per million (TPM). Undetermined values of serum cytokine were imputed with 0. Correlation analysis between serum cytokine protein levels and TPM values in the liver or colon were performed using cor.test function and visualized using ggplot2 package (v4.0.2) in R.

### Quantitative PCR

Purified total RNA was used for qPCR (See method: RNA preparation from colonic epithelial cells and liver). qPCR was performed using Luna Universal One-Step RT-qPCR Kit (NEB, E3005L) according to the manufacture’s protocol. qPCR probes for target genes are listed in Table 2.

**Table 2.**
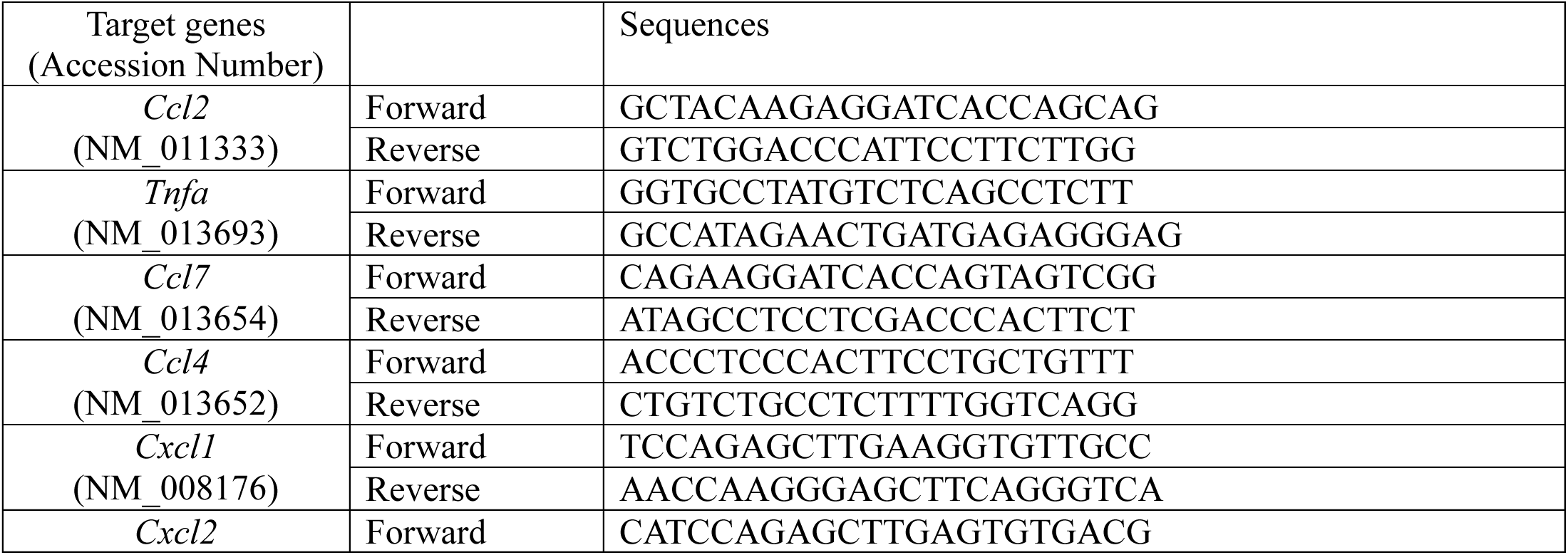
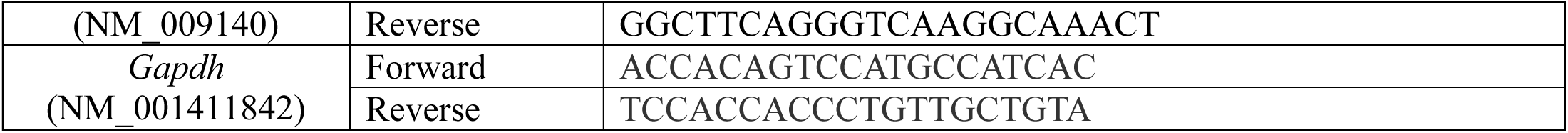
qPCR primers.

### Statistical methods

Statistical analyses were performed using GraphPad Prism version 10. Information regarding the number of samples and statistical tests are described in the figure legends.

## Data availability

The processed RNA-seq data generated in this study have been deposited in the NCBI Gene Expression Omnibus (GEO) under accession number GSE326680. All raw sequences have been deposited in the NCBI Sequence Read Archive (SRA) under accession number PRJNA1445395. The number of differentially expressed genes in KEGG pathways with *p*-values < 0.05 at each time point is listed in Table_Enrich_KEGG_DEG_number.xlsx in Zenodo (https://zenodo.org/records/21264234).

## Code availability

Codes used for data analysis has been deposited and is available through Zenodo (https://zenodo.org/records/21264234).

## Acknowledgements

We thank members of the Waldor lab for helpful discussions and feedback on the manuscript. We thank Elizabeth Scholl for helping us create *C. rodentium* Δ*espF* mutant strain. Funding included Howard Hughes Medical Institute (HHMI) and National Institutes of Health (NIH) R01 AI042347 (M.K.W). The funders had no role in the study design, data collection and analysis, decision to publish, or preparation of the manuscript.

## Author contribution

Conceptualization: Y.H., A.O., M.S., I.W.C., M.K.W.

Data Curation: Y.H., A.O., M.S., T.Y.

Formal Analysis: Y.H., A.O., M.S., T.Y.

Funding Acquisition: M.K.W.

Investigation: Y.H., A.O., M.S., T.Y.

Methodology: Y.H., A.O., M.S.

Project Administration: M.K.W.

Resources: M.K.W.

Software: Y.H.

Supervision: M.K.W.

Validation: Y.H., A.O., M.S.

Visualization: Y.H., A.O., M.S., I.W.C.

Writing – Original Draft Preparation: Y.H., A.O., M.S., I.W.C., M.K.W.

Writing – review & editing: Y.H., A.O., M.S., T.Y., I.W.C., M.K.W.

## License information

This manuscript is subject to the Howard Hughes Medical Institute (HHMI)’s Open Access to Publications policy. HHMI laboratory heads have previously granted the public a non-exclusive CC BY 4.0 license and HHMI a sublicensable license for their research articles. Pursuant to those licenses, the author-accepted manuscript of this manuscript can be made freely available under a CC BY 4.0 license immediately upon publication. This manuscript is the result of funding in whole or in part by the National Institutes of Health (NIH). It is subject to the NIH Public Access Policy. Through acceptance of this federal funding, the NIH has been given the right to make this manuscript publicly available in PubMed Central upon the Official Date of Publication, as defined by the NIH.

## Supporting Information

**Fig. S1 C. rodentium immediately colonizes the cecum following inoculation.** Extended data from Figure 1a. Colony-forming unit (CFU) in the cecum following inoculation with 5×10^9^ CFU. Circles represent data from individual animals, and the solid line represents the geometric mean. The open circle indicates that no bacteria were detected.

**Fig. S2 Principal component scores of transcriptome.** Extended data from Figure 1. **AB.** Top 10 PC scores in the liver (A) or colon (B) from RNA-seq analysis are shown. **CD.** Scree plot showing the fractional variance explained by each principal components (bar graph) for liver**(C)** or colonic epithelium (D) from RNA-seq analysis, and cumulative variance (line graph).

**Fig. S3 Clusters of gene expression patterns in the liver and colonic epithelium during *C. rodentium* infection A. B.** Heatmaps representing the z-score of genes differentially expressed over the course of the study in the liver (A) and colon (B). Differentially expressed genes compared to day 0, with the cut-off (log_2_ fold change > 1, adjusted *p*-value < 0.05) at least one time point, were included. The color code on the y-axis indicates groups identified by k-medoids clustering. The mouse ID on top matches the mouse ID in Fig. 1C and 1F. The panels on the right show the average z-score for genes within each cluster for individual animal at each time point. The symbols correspond to individual mouse IDs shown in the Fig. 1C and 1F.

**Fig. S4 Immunoprofiling in the liver, colonic lamina propria and epithelium AB.** Flow cytometry plot of myeloid cells (A) and lymphocytes (B) analysis in the liver. **CD.** Flow cytometry plot of myeloid cells (C) and lymphocytes (D) analysis in the colonic LP. **EF.** Flow cytometry plot of myeloid cells (E) and lymphocytes (F) analysis in the colonic IEL. **GH**. Flow cytometry plot of Memory CD4+ (G) and CD8+ (H) T cells in IEL fraction (top panels) and the line chart showing the fold changes compared to time-matched uninfected controls (bottom panels).

**Fig. S5 PC1 gene expression in the liver in WT and Δeae strain-infected animals** Z-score of liver RNA expression of top 100 PC1 genes in uninfected control, wt-infected (*wt*), and Δ*eae*-infected (Δ*eae*) animals are shown in color. Uninfected control datasets and four wt-infected samples originate form time series transcriptome dataset. Z-score conversion was performed after data integration, batch correction, and normalization, as described in the Materials and methods.

**Fig. S6 Liver cfu of Δ*espF* and wt *C. rodentium* strain 3 dpi** Wt or Δ*espF C. rodentium* burden (CFU/g) in the liver on 3 days post inoculation. One-way ANOVA after lognormal conversion, and post-hoc test with Holm-Šídák’s multiple comparisons.

**Table S1. Differentially expressed genes in the liver and colon compared to day 0**

**Table S2. Results of pathway analysis with KEGG and MSigDB in the liver and colon**

