## Supplementary Information for "Transcriptomic timeseries links hepatic gene expression to an early and self-limited systemic response to enteric infection"

**Supporting Information**

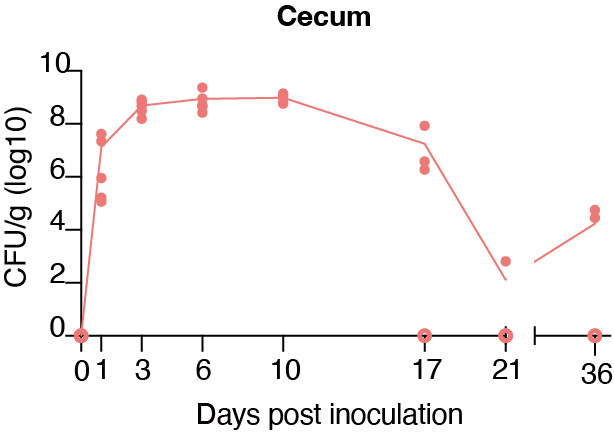

**Fig. S1 *C. rodentium* immediately colonizes the cecum following inoculation.**

Extended data from Figure 1a. Colony-forming unit (CFU) in the cecum following inoculation with 5x10^9^ CFU. Circles represent data from individual animals, and the solid line represents the geometric mean. The open circle indicates that no bacteria were detected.

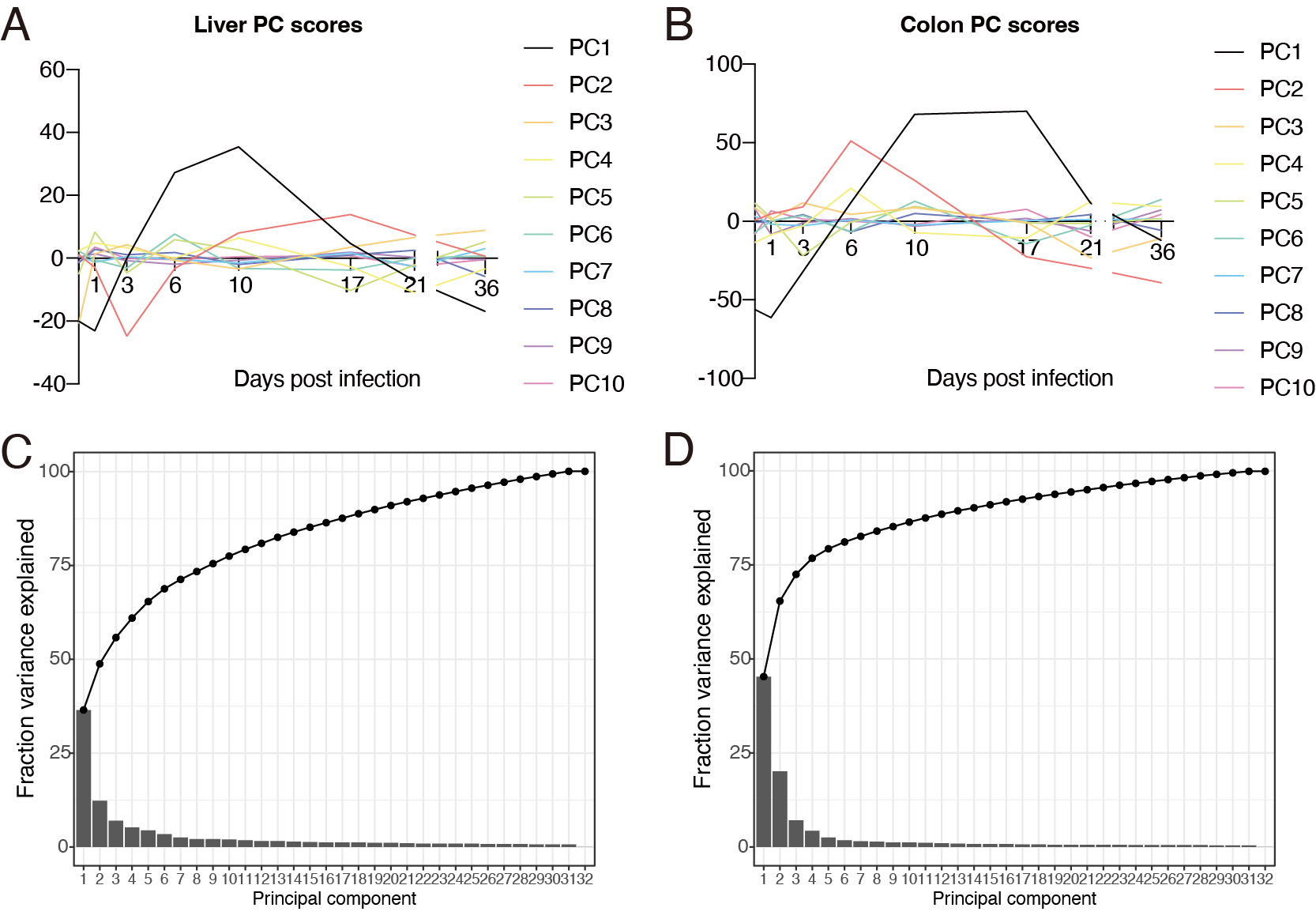

**Fig. S2 Principal component scores of transcriptome.**

Extended data from Figure 1.

**AB.** Top 10 PC scores in the liver (A) or colon (B) from RNA-seq analysis are shown.

**CD.** Scree plot showing the fractional variance explained by each principal components (bar graph) for liver (C) or colonic epithelium (D) from RNA-seq analysis, and cumulative variance (line graph).

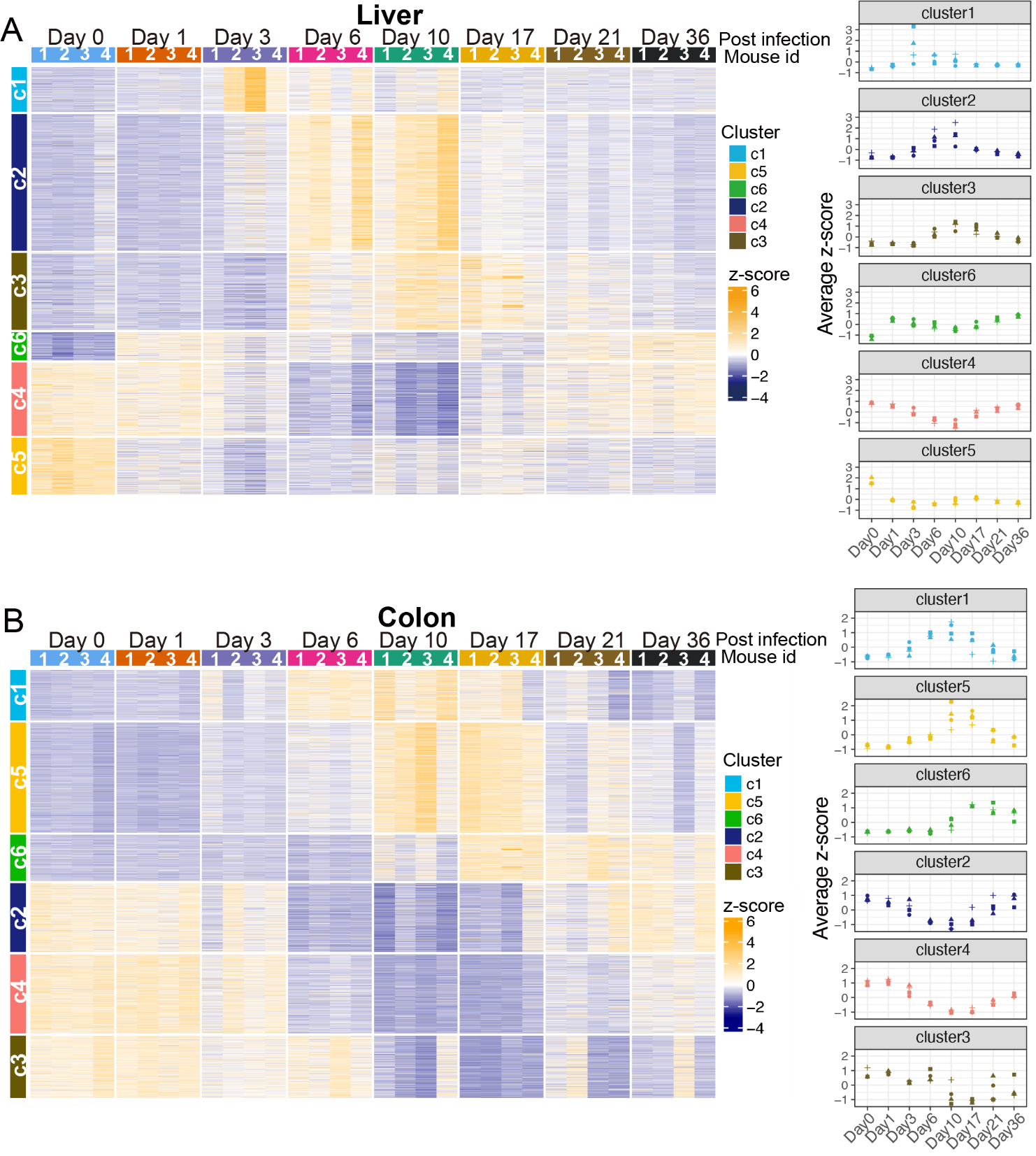

**Fig. S3 Clusters of gene expression patterns in the liver and colonic epithelium during *C. rodentium* infection**

**A. B.** Heatmaps representing the z-score of genes differentially expressed over the course of the study in the liver (A) and colon (B). Differentially expressed genes compared to day 0, with the cut-off (log_2_ fold change > 1, adjusted p-value < 0.05) at least one time point, were included. The color code on the y-axis indicates groups identified by k-medoids clustering **The mouse ID on top matches the mouse ID in Fig. 1C and 1F. The panels on the right show the average z-score for genes within each cluster for individual animal at each time point. The symbols correspond to individual mouse IDs shown in the Fig. 1C and 1F.**

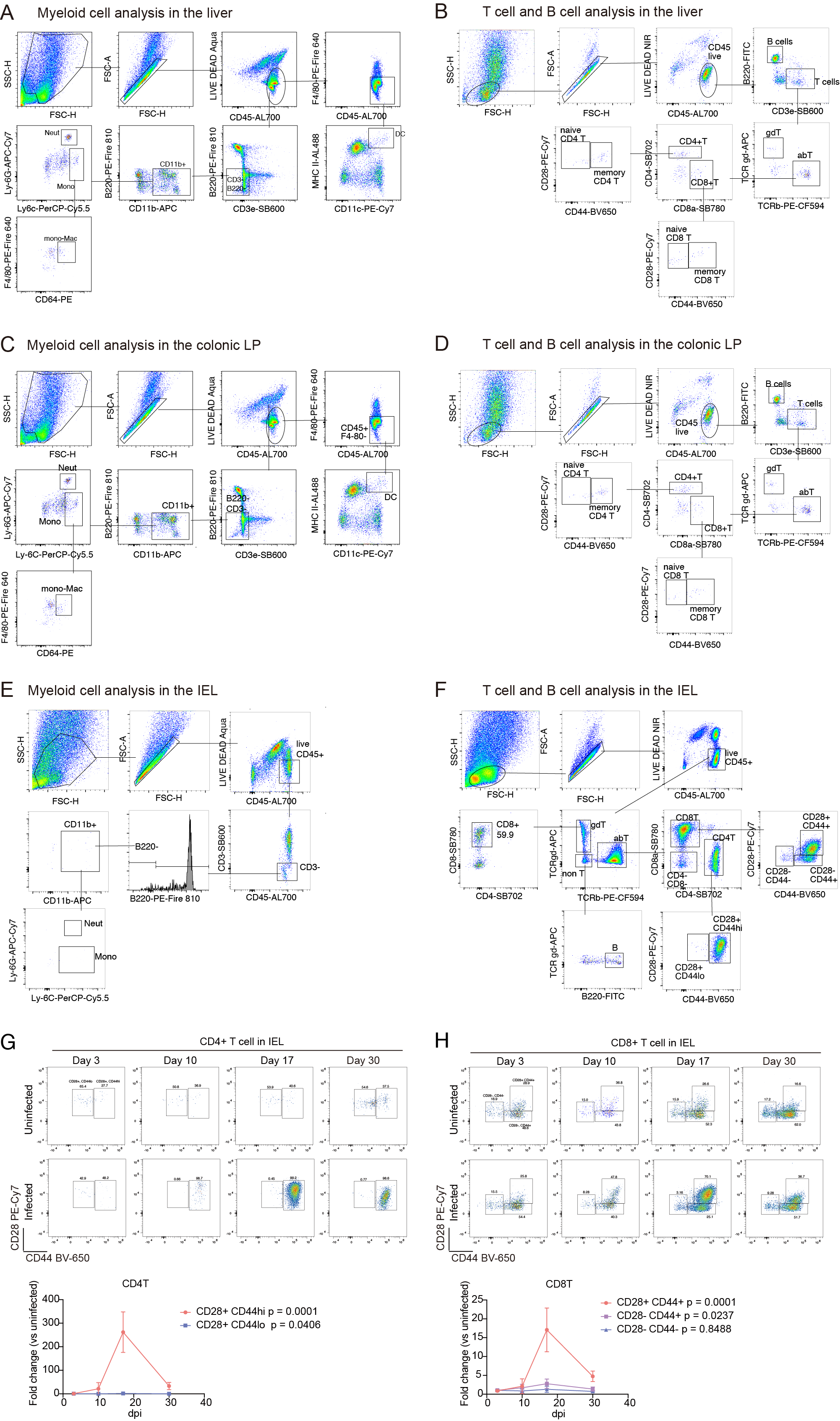

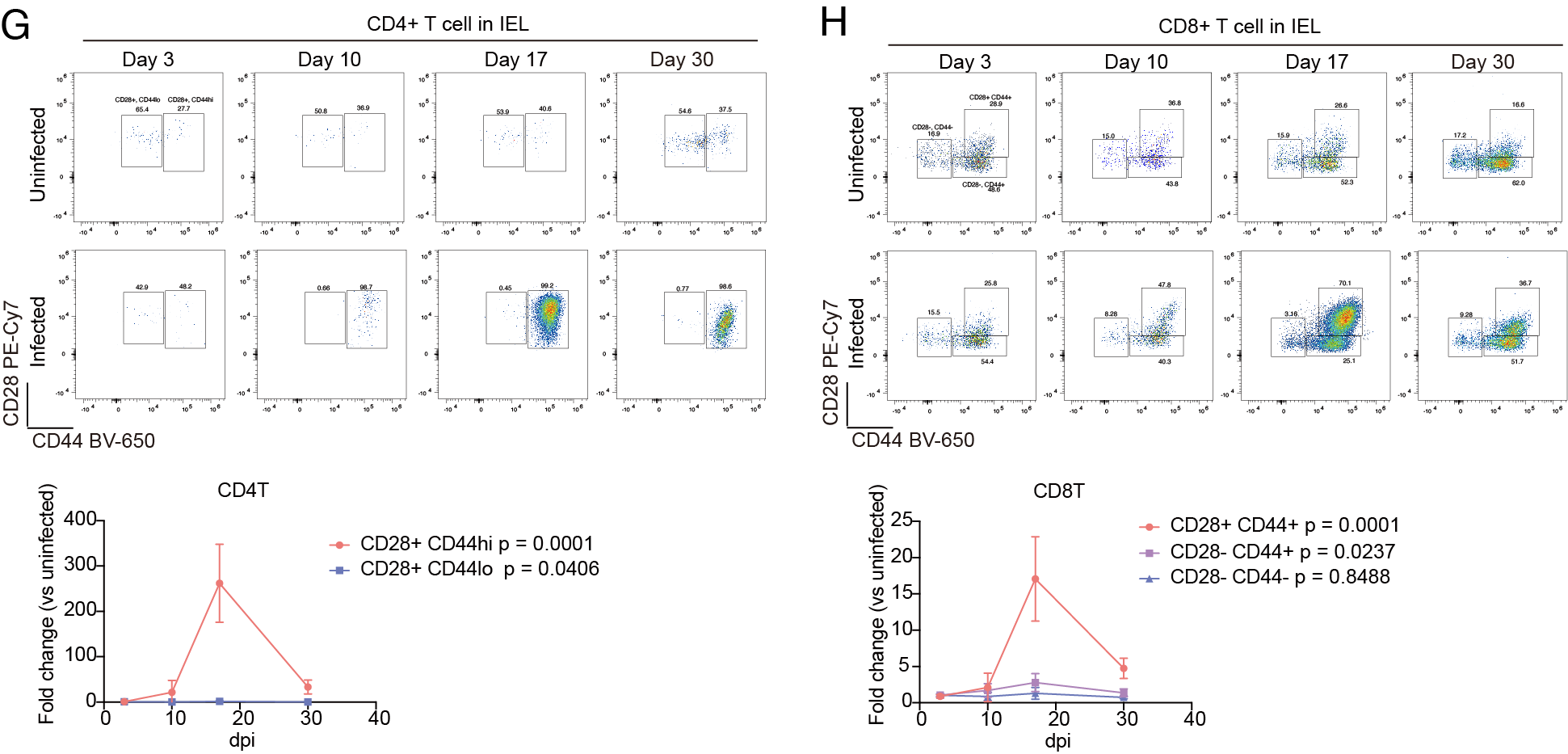

**Fig. S4 Immunoprofiling in the liver, colonic lamina propria and epithelium**

**AB.** Flow cytometry plot of myeloid cells (A) and lymphocytes (B) analysis in the liver.

**CD.** Flow cytometry plot of myeloid cells (C) and lymphocytes (D) analysis in the colonic LP.

**EF.** Flow cytometry plot of myeloid cells (E) and lymphocytes (F) analysis in the colonic IEL.

**GH**. Flow cytometry plot of Memory CD4+ (G) and CD8+ (H) T cells in IEL fraction (top panels) and the line chart showing the fold changes compared to time-matched uninfected controls (bottom panels).

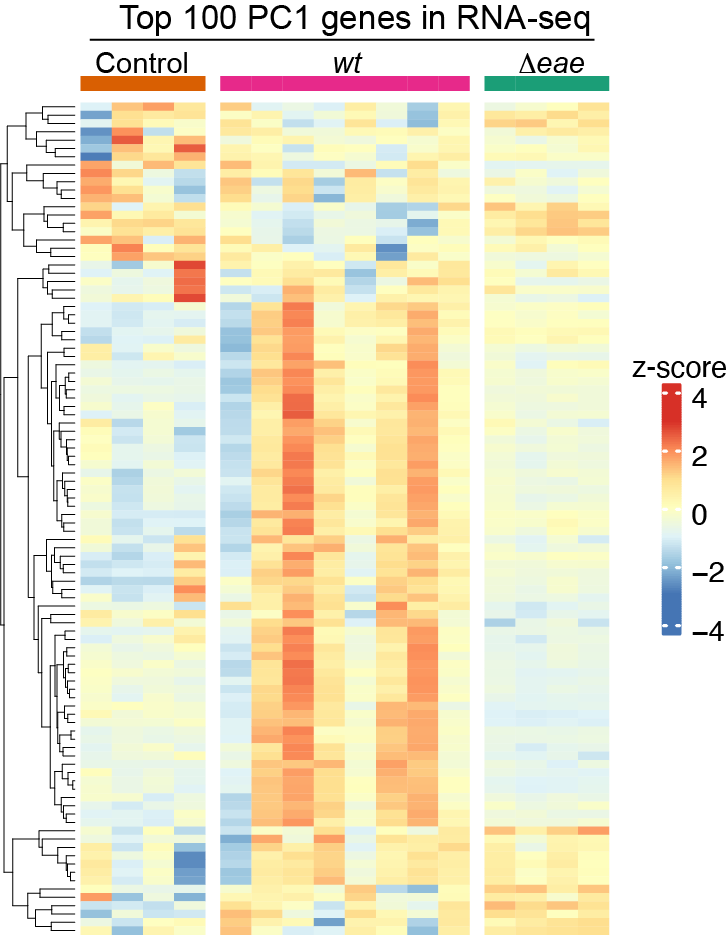

**Fig. S5 PC1 gene expression in the liver in WT and Δ*eae* strain-infected animals**

Z-score of liver RNA expression of top 100 PC1 genes in uninfected control, wt-infected (*wt*), and Δ*eae*-infected *(*Δ*eae*) animals are shown in color. Uninfected control datasets and four wt-infected samples originate form time series transcriptome dataset. Z-score conversion was performed after data integration, batch correction, and normalization, as described in the Materials and methods.

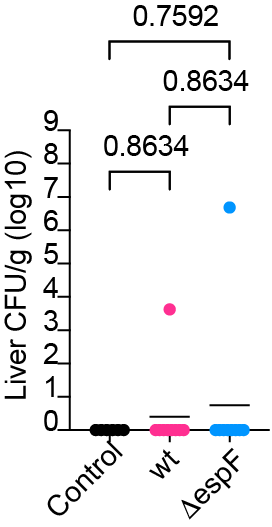

**Fig. S6 Liver cfu of Δ*espF* and wt *C. rodentium* strain 3 dpi**

Wt or Δ*espF C. rodentium* burden (CFU/g) in the liver on 3 days post inoculation. One-way ANOVA after lognormal conversion, and post-hoc test with Holm-Šídák's multiple comparisons.
